# Localization of immunoreactive, dystonia-associated protein torsinA near the Golgi apparatus of cultured rodent astrocytes

**DOI:** 10.1101/2019.12.11.873554

**Authors:** Sadahiro Iwabuchi, Hiroyuki Kawano, N. Charles Harata

## Abstract

An in-frame deletion of a single glutamic acid codon in the *TOR1A* gene causes the neurological disorder DYT1 dystonia, but the cellular pathophysiology of this disorder remains elusive. A current model postulates that the wild-type (WT) torsinA protein is mainly localized to the endoplasmic reticulum (ER), but that the mutant form (ΔE-torsinA) is diverted to the nuclear envelope and cytoplasmic inclusion bodies. This mis-localization has been observed by overexpressing the proteins in neuronal and non-neuronal cells. However, it is not clear whether this model is valid for the astrocytic glial cells that support and modify neuronal functions. Here we report, using rodent astrocytes in primary culture, that the overexpressed torsinA proteins were distributed as predicted by the mis-localization model. We also found by immunocytochemistry that the cultured astrocytes express torsinA endogenously. Most of the signals from endogenous protein, whether the WT or ΔE form, were localized near a *cis*-Golgi marker GM130. Such localization of endogenous proteins was found in glial cells from several sources: the hippocampus of WT rats, the hippocampus and striatum of WT mice, and the hippocampus and striatum of ΔE-torsinA knock-in mice, a model of DYT1 dystonia. These results show that the mis-localization model is applicable to overexpressed torsinA proteins, but not applicable to those expressed at endogenous levels, at least in cultured rodent astrocytes. These discrepancies in the distribution of overexpressed versus endogenous torsinA proteins highlight the potential need for caution in interpreting the results of overexpression studies.

## 1. INTRODUCTION

Mutation of the *TOR1A* gene is associated with DYT1 dystonia (Ozelius et al., 1997), an inherited, autosomal-dominant movement disorder that is clinically defined as “early-onset generalized isolated dystonia” (Albanese et al., 2013). The pathogenic mutation (c.904_906delGAG/c.907_909delGAG; p.Glu302del/p.Glu303del) leads to an in-frame deletion of glutamic acid residue from the encoded protein, torsinA (ΔE-torsinA). TorsinA belongs to the <u>A</u>TPases <u>a</u>ssociated with diverse cellular <u>a</u>ctivities (AAA+) family of proteins, whose members generally perform chaperone-like functions, assisting in protein unfolding, protein-complex disassembly, membrane trafficking, and vesicle fusion (Hanson and Whiteheart, 2005; White and Lauring, 2007; Burdette et al., 2010; Zhao et al., 2013). The lack of neurodegeneration in the patients’ brains (Standaert, 2011) suggests that the abnormality that underlies DYT1 dystonia affects neural connectivity and neuronal functions (Breakefield et al., 2008), yet the details of the pathophysiological mechanisms remain unclear.

According to one model, a part of the cellular basis of DYT1 dystonia pathophysiology is the mis-localization of ΔE-torsinA protein, as reviewed e.g. in (Granata et al., 2009). The wild-type (WT) torsinA protein has been reported to be distributed diffusely throughout the cytoplasm and co-localized with the endoplasmic reticulum (ER), with a minor component in the contiguous nuclear envelope. The ΔE-torsinA protein is concentrated on the nuclear envelope where it induces structural abnormalities, or localized in cytoplasmic inclusion bodies. The supporting evidence comes from studies in which human torsinA proteins were overexpressed in many types of non-neuronal cells (Kustedjo et al., 2000), in addition to neurons (Nery et al., 2011) and neural tumor cells (Hewett et al., 2000; Gonzalez-Alegre and Paulson, 2004; Misbahuddin et al., 2005; Gordon and Gonzalez-Alegre, 2008), as reviewed in (Harata, 2014). Such mutation-associated mis-localization has been hypothesized to constitute an important pathogenic event in DYT1 dystonia.

Astrocytic glial cells comprise the most abundant cell population in the brain. These cells support neurons by maintaining the local environment, and also actively regulate neuronal functions such as network formation (Molofsky et al., 2014), and synaptic transmission in the striatum (Goubard et al., 2011) and hippocampus (Panatier et al., 2011). They are also involved in pathological processes and the protective mechanisms triggered by cellular insults (Nedergaard et al., 2010; Allaman et al., 2011; Iwabuchi et al., 2014a). Thus a change in the condition of these cells can lead to neuronal dysfunction. Nevertheless, the mis-localization model of ΔE-torsinA has not been tested in astrocytes of mammalian brains. Human WT-torsinA was overexpressed in human astrocytes in primary culture, and was found to be distributed diffusely in the cytoplasm and co-localized with an astrocyte marker, the cytoskeletal, intermediate-filament protein glial fibrillary acidic protein (GFAP), in both the cell body and the processes (Armata et al., 2008). While this result is consistent with the assumed localization of WT-torsinA to the ER, ΔE-torsinA has not been overexpressed in astrocytes. Thus the validity of the mis-localization of ΔE-torsinA in this specific cell type remains to be assessed.

At least three further pieces of information require clarification, regarding the subcellular distribution of torsinA in astrocytes. The first is whether glial cells express the endogenous torsinA. TorsinA protein has been detected in glial cells of the mouse brain (Kim et al., 2010; Puglisi et al., 2013). However, these findings are inconsistent with other reports, including analyses of human brains (Shashidharan et al., 2000; Rostasy et al., 2003) and mouse brains (Martella et al., 2009). If torsinA is expressed in glial cells, it is unclear why it has been overlooked in these studies. Second, if torsinA is expressed in glial cells, which subcellular compartments is it located in? This is a critical question because the distribution of exogenously introduced, overexpressed protein may not reflect the endogenous state. The subcellular distribution of torsinA protein at endogenous level has not been examined in detail in glial cells. Notably, patients with DYT1 dystonia are heterozygous for the ΔE-torsinA mutation (*TOR1A*^+/ΔE^), and the total level of torsinA expression is either equivalent to or slightly less than that in normal human subjects – but not higher (Goodchild and Dauer, 2004; Goodchild et al., 2005). The third question is to what extent the subcellular localization of endogenous torsinA varies across astrocytes, which are a highly heterogenous cell population. For example, astrocytes in different brain regions (Tsai et al., 2012; Molofsky et al., 2014) and different animal species (Ahlemeyer et al., 2013) have been reported to differ both functionally and morphologically.

Here we have addressed the following questions about torsinA expression: 1) whether its overexpression in astrocytes changes its subcellular distribution from a diffuse ER-like pattern of WT-torsinA to the staining of the nuclear envelope and inclusion bodies by ΔE-torsinA; 2) whether endogenous expression is detectable in astrocytes; 3) whether endogenous expression of ΔE-torsinA differs from that of WT-torsinA; and 4) whether the endogenous distribution in astrocytes varies by brain region and by rodent species. For addressing these questions, astrocytes were obtained from WT rats and the ΔE-torsinA knock-in mouse model of DYT1 dystonia (Dang et al., 2005; Goodchild et al., 2005). Heterozygous ΔE-torsinA knock-in mice (HET, *Tor1a*^+/ΔE^) are viable and geno-copy the DYT1 dystonia patients in that they carry each of the WT and mutant allele. In these mice, total torsinA is not upregulated, whether at the mRNA or protein level (Goodchild et al., 2005; Yokoi et al., 2010). Primary cultures of astrocytes were used, because these large and flat cells allow imaging protein localization at high spatial resolution. They were prepared from the hippocampus and striatum, the brain regions with high expression of torsinA protein (Walker et al., 2001).

## 2. EXPERIMENTAL PROCEDURES

### 2.1. Genotyping and animals

All animal care and procedures were approved by the University of Iowa Animal Care and Use Committee, and were performed in accordance with the standards set by the National Institutes of Health Guide for the Care and Use of Laboratory Animals (NIH Publications No. 80-23, revised 1996). Every effort was made to minimize suffering of the animals. On postnatal days 0-1, pups of the ΔE-torsinA knock-in mouse model of DYT1 dystonia (Goodchild et al., 2005), of both sexes, were genotyped using a fast-genotyping procedure (EZ Fast Tissue/Tail PCR Genotyping Kit, EZ BioResearch LLC, St, Louis, MO) (Koh et al., 2015). Newborn pups of both sexes were obtained from Crl:CD(SD) (Sprague-Dawley) rat mothers (Charles River Laboratories International, Wilmington, MA).

### 2.2. Cell culture

Glial cells from individual newborn mouse pups were cultured using a modification of our previously reported protocol (Koh et al., 2013; Koh et al., 2015). In brief, the striatum and hippocampus were dissected on postnatal days 0-1. The hippocampal preparation included the CA3-CA1 region, but excluded the dentate gyrus (Harata et al., 2006; Kakazu et al., 2012). The dissected tissues were trypsinized, mechanically dissociated, plated in an uncoated T25 culture flask and cultured in a humidified incubator (5% CO_2_-95% O_2_, 37°C). At 7-10 days *in vitro* (DIV), the glial cells were trypsinized and plated on 12-mm glass coverslips (thickness No. 0, Carolina Biological Supply, Burlington, NC) previously coated with Matrigel Basement Membrane Matrix solution (BD Biosciences, San Jose, CA). To avoid the overlap of signal from multiple glial cells, we plated the cells at a low density, i.e., at ~10^4^ live cells / coverslip. 2-3 days after plating on coverslips, the cells were treated with 81 µM 5-fluoro-2’-deoxyuridine and 205 µM uridine, to inhibit cellular proliferation. They were used for experiments 19-24 days after plating on coverslips.

Glial cells from the hippocampus of individual newborn rat pups were cultured in much the same way as mouse cells, but with the following modifications. The dissected tissues were trypsinized, mechanically dissociated and plated in a T25 culture flask coated with Matrigel. After being cultured in the incubator for 2-3 days, non-adherent or weakly adherent cells were removed. At 6-8 DIV, the glial cells were trypsinized and plated in an un-coated T75 flask, and further cultured in the incubator. At 18-20 DIV, the glial cells were trypsinized and plated on Matrigel-coated coverslips. The plating density was ~10^3^ live cells / coverslip. 3-4 days after plating on coverslips, the cells were treated with 2 µM cytosine β-D-arabinofuranoside to inhibit proliferation. They were used for experiments 19-24 days after plating on coverslips.

Neuron-glia co-cultures were prepared as described previously (Koh et al., 2013; Koh et al., 2015) from the hippocampus of WT rats.

In all experiments, data were obtained from 2-6 separate culture batches (i.e., pups).

### 2.3. Immunocytochemistry

Our published protocol (Koh et al., 2013; Mitchell et al., 2018) was used with slight modifications. The cultured cells were fixed with 4% paraformaldehyde (15710, Electron Microscopy Sciences, Hatfield, PA) and 4% sucrose in Tyrode’s solution (containing, in mM: 125 ^NaCl, 2 KCl, 2 CaCl^2^, 2 MgCl^2^, 30 D-glucose, 25 HEPES, pH 7.4 adjusted with 5 M NaOH, ~310^ mOsm without adjustment) for 30 min at 4°C. The cells were washed with Tyrode’s solution, twice for 5 min per wash at 4°C. They were permeabilized with 0.1% Triton X-100 in Tyrode’s solution for 10 min, and washed with Tyrode’s solution three times (5 min per wash). They were blocked with 2% normal goat serum (G-9023, Sigma-Aldrich, St. Louis, MO) in phosphate-buffered saline (PBS, 70011-044, pH 7.4, GIBCO-Life Technologies) (blocking solution), for 60 min at room temperature. Thereafter they were treated with the following primary antibodies (diluted in blocking solution) overnight (15-21 hours) at 4°C.

Rabbit polyclonal antibodies were: anti-torsinA (ab34540, Abcam, Cambridge, MA; 400x dilution), and anti-glucose-regulated protein of 78 kDa / binding immunoglobulin protein (GRP78/BiP) (PA1-014A, Thermo Fisher Scientific, Waltham, MA; 4000x dilution). Mouse monoclonal antibodies were: anti-Golgi matrix protein of 130 kDa (GM130) (610822, BD Biosciences, San Jose, CA; 400x dilution), anti-protein disulfide isomerase (PDI) (ADI-SPA-891, Enzo Life Sciences / StressGen, Farmingdale, NY; 400x dilution), anti-nuclear pore complex (NPC) proteins (ab24609, Abcam; 400x dilution), anti-GFAP monoclonal cocktail (NE1015, Merck Millipore, Billerica, MA; 1000x dilution), and anti-green fluorescent protein (GFP) (DSHB-GFP-12A6, Developmental Studies Hybridoma Bank, Iowa City, IA; 1000x dilution).

Following washing with PBS 3 times (7 min per wash), the cells were incubated with goat anti-rabbit or anti-mouse IgG antibody conjugated with Alexa Fluor 488 dye or Alexa Fluor 568 dye (Life Technologies; 1000x dilution in blocking solution) for 60 min at room temperature. They were washed with PBS at least 5 times (20 min per wash), transferred to an imaging chamber (RC-26, Warner Instruments, Hamden, CT), and observed directly in PBS.

In some experiments, the nuclei were counterstained using membrane-permeant Hoechst 33342 (Life Technologies). After the final wash in immunocytochemical procedures, the cells were treated with the dye at 0.5 µg/ml in PBS for 10 min at room temperature, and washed for 3-5 min with PBS. The cells were then transferred to the imaging chamber.

### 2.4. TorsinA overexpression

Cultured rat hippocampal glial cells were transfected 14-16 days after plating on coverslips, with a plasmid construct encoding either the human WT- or ΔE-torsinA tagged with GFP (Naismith et al., 2004). The calcium phosphate-based CalPhos Mammalian Transfection Kit (631312, Clontech Laboratories, Mountain View, CA) was used (Harata et al., 2006), with a modified procedure for mixing solutions (Jiang and Chen, 2006). Briefly, transfection was carried out in 24-well culture plates, using 2 μg DNA per well in Minimum Essential Medium (MEM) (51200-038, GIBCO-Life Technologies, Grand Island, NY). Cells were exposed to the precipitate at 37°C for 40-70 min, washed three times with MEM, and returned to the original culture medium. Transfected cells were observed 7 days after transfection (or ~22 days after plating on coverslips), live and without fixation.

### 2.5. Cell imaging

Imaging was confined to glial cells that formed confluent or nearly confluent cultures, as judged by differential interference contrast (DIC) microscopy or phase-contrast microscopy. The selected glial cells did not display signs of deterioration, such as the formation of large, spherical intracellular vacuoles. For quantitative analysis, imaging was restricted to those glial cells that had nuclei that were easily identified by fluorescence or transmitted light microscopy, and that formed a monolayer where ER or Golgi structures of neighboring cells did not overlap. All imaging experiments were performed at room temperature (23-25°C).

### 2.6. Fluorescence imaging system

For general characterization by widefield optics, the cells were imaged using an inverted microscope (Eclipse-TiE, Nikon, Melville, NY) equipped with an interline CCD camera (Clara, Andor Technology, Belfast, UK). The camera was cooled at −45°C by an internal fan. Alexa Fluor 488 dye was excited using a 490-nm light-emitting diode (LED, CoolLED-Custom Interconnect, Hampshire, UK) and imaged with a filter cube (490/20-nm excitation, 510-nm dichroic long-pass, 530/40-nm emission). Alexa Fluor 568 dye was excited using a 595-nm LED (CoolLED-Custom Interconnect) and imaged with a filter cube (590/55-nm excitation, 625-nm dichroic long-pass, 665/65-nm emission). The LED was used at 100% intensity, and exposure time was 1 sec. Hoechst 33342 was excited using a 400-nm LED (CoolLED-Custom Interconnect) at 100% intensity, and imaged with a filter cube (405/40-nm excitation, 440-nm dichroic long-pass, 470/40-nm emission), for a 0.2-sec exposure. 16-bit images were acquired using a 40x objective lens (Plan Fluor, numerical aperture 1.30, Nikon) with a coupler (0.7x) and without binning, in the single-image capture mode of the Solis software (Andor).

For quantitative characterization, the cells were imaged using a confocal laser-scanning microscope (510 confocal, Carl Zeiss MicroImaging GmbH, Göttingen, Germany) equipped with an inverted microscope (Axiovert 100M, Carl Zeiss) and a 40x objective lens (Plan-Neofluor, numerical aperture 1.30). Alexa Fluor 488 dye was excited using the 488-nm line of an argon laser at 1% intensity, and imaged using a 505-530 nm band-pass emission filter. Alexa Fluor 568 dye was excited using 543-nm line of helium-neon laser at 100% intensity, and imaged using a 585 nm long-pass emission filter. Voxel sizes were 0.44 x 0.44 x 0.40 µm or 0.22 x 0.22 x 0.40 µm (x-y-z). Pinhole size was 58 µm with an Airy unit of 0.8. Pixel time was 2.51 µs. Four frames were averaged to give a single 12-bit image at each level of focus (optical section), and a stack of images (z-stack) was acquired at focusing levels 0.40-µm apart.

Imaging conditions were kept the same for each brain region, such that the absolute intensities can be compared across cells.

### 2.7. Image analysis

The images were opened in a 16-bit format and analyzed using the ImageJ software (W. S. Rasband, National Institutes of Health, Bethesda, MD).

For quantifying the intensity of immunoreactive torsinA staining in glial cells, the effect of noise was minimized by averaging three consecutive images in a confocal z-stack and filtering with the Gaussian-blur function. The radius of the Gaussian filter was 1.0 µm with pixel sizes of 0.44 x 0.44 µm or 0.5 µm with pixel sizes of 0.22 x 0.22 µm. For measuring the torsinA intensity in the Golgi apparatus, regions of interest (ROI’s) on the Golgi apparatus were assigned using the GM130 channel, with the intensity threshold set as the mean intensity + 3x standard deviation in a negative control image (primary antibody omission) that was acquired under the same imaging condition. These ROI’s were transferred to the torsinA channel, and the pixel intensities in the ROI’s were averaged to give a single value for one cell. For measuring the torsinA intensity in the ER and nucleoplasm, ROI’s were manually assigned in the PDI channel. The ER region excluded the Golgi apparatus, and the nucleoplasm excluded the nucleoli. These ROI’s were transferred to the torsinA channel and the averaged pixel intensities provided single values for one cell. In this study, the number of cells was counted as the number of nuclei.

For comparing the torsinA intensity in glial cells and neurons in the same image field, the images in a z-stack were projected to a single image using the maximum intensity projection (MIP) algorithm. TorsinA intensity in the Golgi apparatus was measured as described above.

For presenting images, they were filtered using the Gaussian-blur function. They were then converted from 16-bit to 8-bit format, with the minimal and maximal intensity in individual images assigned values of 0 and 255, respectively, except where otherwise mentioned. Where intensity is compared under the same imaging conditions, the intensity range was determined using the whole image series for comparison, and converted using the same intensity scale.

### 2.8. Characterization of antibodies used in the current study

Anti-torsinA antibody used in this study is a rabbit polyclonal antibody. The specificity of this antibody was demonstrated in negative controls, by immunohistochemical staining of the brains of conditional *Tor1a* knock-out mice (Liang et al., 2014), the brains of WT mice after antigen preadsorption (Puglisi et al., 2013), and the cultured rodent brain neurons and glial cells after antigen preadsorption (Koh et al., 2013). The negative control also includes the Western blotting of brain homogenates from *Tor1a* knock-out mice (Kim et al., 2010; Yokoi et al., 2010). This antibody was used for immunohistochemical staining of the brains of WT mice (Puglisi et al., 2013), the brains of transgenic mice overexpressing ΔE-torsinA (Napolitano et al., 2010; Sciamanna et al., 2011), and cultured brain neurons of ΔE-torsinA knock-in mice (Koh et al., 2013). It was also used in Western blotting experiments to detect total torsinA protein (Cao et al., 2010; Kim et al., 2010; Yokoi et al., 2010; Gordon et al., 2011; Puglisi et al., 2013; Yokoi et al., 2013). Similarly to other anti-torsinA antibodies (Goodchild et al., 2005), it detects both WT- and ΔE-torsinA proteins (Cao et al., 2010; Napolitano et al., 2010; Sciamanna et al., 2011; Koh et al., 2013).

Anti-GM130 is a mouse monoclonal antibody that recognizes a Golgi marker. The immunocytochemical labeling pattern in the current study was the same as that observed in cultured astrocytes isolated from the mouse cerebral cortex (Ramamoorthy and Whim, 2008). Similar staining was observed in cultured cells other than astrocytes; they include the human adenocarcinoma line HeLa cells (Roy et al., 2012; Simpson et al., 2012), rat hippocampal neurons (Horton and Ehlers, 2003; Horton et al., 2005), and mouse cerebral cortex neurons (Vitry et al., 2010). It was also observed in neurons *in vivo*, in the rat hippocampus (Horton et al., 2005; Cox and Racca, 2013), the rat cerebral cortex (Horton et al., 2005), the mouse hippocampus (Clarke et al., 2009) and the mouse cerebral cortex (Condon et al., 2013).

Anti-PDI is a mouse monoclonal antibody that recognizes an ER marker. This antibody was used for immunocytochemical staining of human brains (Walker et al., 2002), HeLa cells (Chadrin et al., 2010) and cultured mouse fibroblasts (Kornak et al., 2001).

Anti-GRP78/BiP is a rabbit polyclonal antibody that recognizes an ER marker. This antibody has been used for immunocytochemical staining of rat brain gliosarcoma 9L cells, where it co-localized with PDI (Sun et al., 2006), as well as of the human retinal pigment epithelium (Marmorstein et al., 2000).

Anti-NPC is a mouse monoclonal antibody that recognizes a nuclear envelope marker. This antibody has been used for immunocytochemical staining of the mouse hippocampal line HT22 cells (Benz et al., 2010), HeLa cells (Umlauf et al., 2013), and Chinese hamster ovary CHO cells (Chen et al., 2011).

Anti-GFAP is a mouse monoclonal antibody that recognizes an astrocyte marker. This antibody has been used for immunocytochemical staining of brains of rat (Casella et al., 2004), mouse (Meikle et al., 2007) and human (McLendon et al., 1986; Desilva et al., 2007).

Anti-GFP is a mouse monoclonal antibody. This antibody has been used for immunocytochemical staining of GFP overexpressed in human skin epidermoid carcinoma A431 cells and the zebrafish embryonic neurons (Sanchez et al., 2014).

As a secondary antibody control, the primary antibodies were omitted from the immunocytochemical procedures in each experiment. No labeling was observed under the same imaging conditions and contrast of acquired images (Koh et al., 2013).

### 2.9. Drugs

All chemical reagents were purchased from Sigma-Aldrich unless otherwise specified.

### 2.10. Statistical analysis

Means were compared using the unpaired Student’s t test. All significance values provided are two-tailed p values.

## 3. RESULTS

### 3.1. Subcellular distribution of overexpressed torsinA proteins in cultured rat glial cells

We first examined the subcellular distribution of torsinA proteins following their overexpression in cultured rat hippocampal glial cells. The cells formed an almost confluent sheet, as shown by DIC imaging (Fig. 1A). Immunocytochemistry revealed that many of these cultured glial cells were positive for the astrocyte marker GFAP (Fig. 1A). As expected, the GFAP-immunoreactive structure was fibrous and was excluded from the nuclei that were identified by Hoechst dye (Fig. 1A). The nuclei occupied only a small area.

**Fig. 1.**
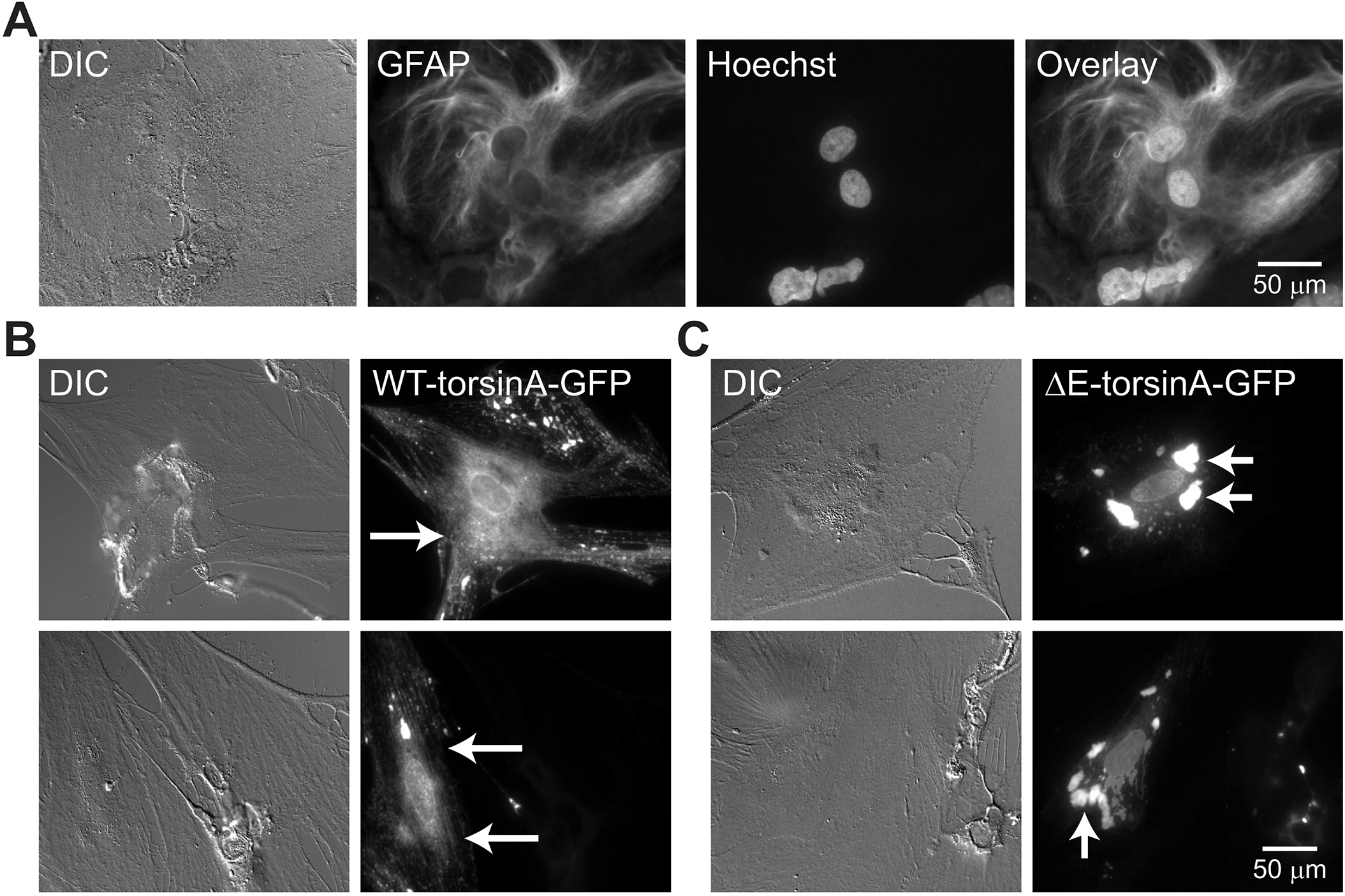
Localization of overexpressed, GFP-tagged wild-type (WT)- and ΔE-torsinA protein in cultured glial cells from the rat hippocampus. Differential interference contrast (DIC) microscopy and widefield fluorescence microscopy images. **A**) Cells co-stained with an antibody against the astrocyte marker glial fibrillary acidic protein (GFAP) and the nucleoplasmic Hoechst dye. GFAP is visualized using Alexa Fluor 488 dye (green channel) and Hoechst dye is visualized in the blue channel. Astrocytes in this field were confluent. **B**) Two representative cells (top and bottom) overexpressing GFP-tagged WT-torsinA (WT-torsinA-GFP). GFP expression (green channel) is diffuse in the cytoplasm (long arrows), and additional fluorescence is observed in the perinuclear region. **C**) Two representative cells overexpressing GFP-tagged ΔE-torsinA (ΔE-torsinA-GFP). GFP expression is in large cytoplasmic inclusion bodies (short arrows). GFP was imaged in live cells.

The cultured rat glial cells were transfected with plasmid constructs encoding GFP-tagged torsinA proteins. The GFP signal for the overexpressed WT-torsinA (WT-torsinA-GFP) was distributed diffusely (long arrows, Fig. 1B), with a minor component surrounding the nuclei (perinuclear). This finding is consistent with results from an earlier study in cultured human astrocytes (Armata et al., 2008).

Overexpressed ΔE-torsinA (ΔE-torsinA-GFP), in contrast, was present mostly in large cytoplasmic inclusion bodies (short arrows, Fig. 1C), consistent with observations in other cell types (Hewett et al., 2000; Kustedjo et al., 2000). These results support the mis-localization model, i.e. that the localization ΔE-torsinA differs from that of WT-torsinA in the overexpression system.

### 3.2. Subcellular localization of endogenous WT-torsinA protein in cultured rat glial cells

Immunocytochemistry was used to evaluate the distribution of endogenous WT-torsinA in cultured hippocampal glial cells from WT rats (Fig. 2). In contrast to the results from the overexpression analysis (Fig. 1B), this experiment did not reveal diffusely distributed torsinA signal (Fig. 2A). Moreover, the strongest intensity was just outside individual nuclei, though not necessarily surrounding the nuclei (paranuclear) ("Hoechst" in Fig. 2A). The intensity of staining was evaluated objectively by measurement along a straight line (bottom panel in Fig. 2A). There was weak cytoplasmic staining. In some regions the intensity was higher than in the cytoplasm. In particular, strong peaks were present near the nucleus. Low-level staining was also detected in the nucleoplasm, with gaps that may represent the nucleoli.

**Fig. 2.**
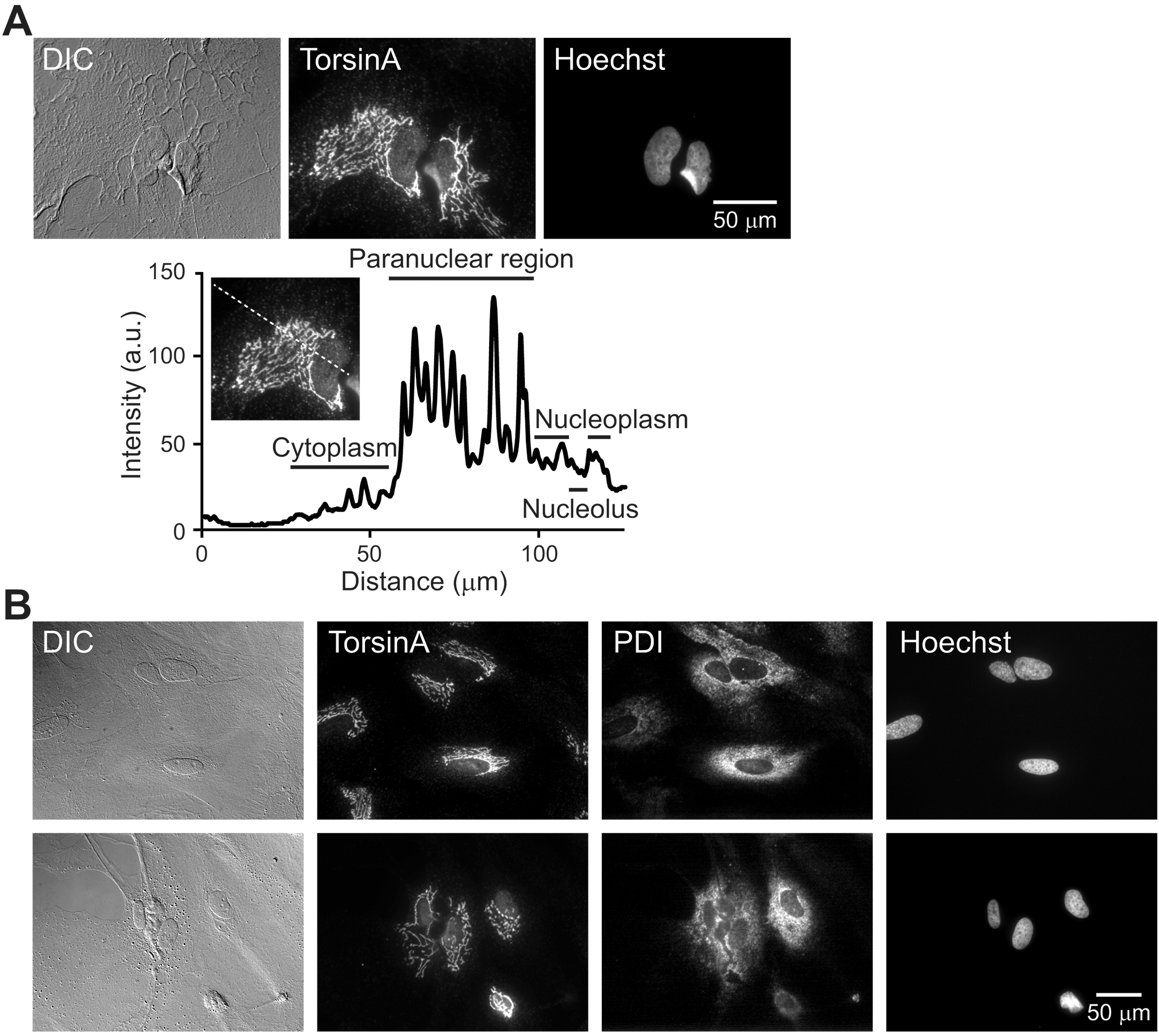
Localization of endogenous WT-torsinA in cultured glial cells from the rat hippocampus. **A**) DIC and widefield fluorescence images of glial cells stained by immunocytochemistry for endogenous torsinA (Alexa Fluor 488) and counterstained with Hoechst dye. Graph in the bottom panel illustrates the fluorescence intensity along the dotted line drawn in the inset. Intensity was measured after subtracting the minimal intensity value in the image field. Immunoreactive torsinA was mainly paranuclear (i.e., near the nucleus but not necessarily surrounding it). Minor expression level was found in the nucleoplasm and cytoplasm. **B**) DIC and widefield fluorescence images of glial cells stained by immunocytochemistry for endogenous torsinA (Alexa Fluor 488) and the endoplasmic reticulum (ER) marker, protein disulfide isomerase (PDI; Alexa Fluor 568). The two distributions are similar but not identical. Two representative fields are shown.

This distribution pattern for torsinA was compared to that of the ER (Fig. 2B) because the overexpressed WT-torsinA in non-astrocytic cells is co-localized with the ER markers PDI (O’Farrell et al., 2002; Bragg et al., 2004b; Naismith et al., 2004; Misbahuddin et al., 2005; Jungwirth et al., 2010; Nery et al., 2011; Hettich et al., 2014), GRP78/BiP (Kustedjo et al., 2000) and KDEL (Goodchild and Dauer, 2004; Calakos et al., 2010; Maric et al., 2011). Double staining revealed that the distributions of torsinA and PDI were similar (Fig. 2B), with immunoreactivity strong near the nuclei and less prominent as distance from the nucleus increased, although DIC shows that almost all parts of the image field contained glial cells (Fig. 2B). However, the distributions were not identical. PDI was more diffuse than torsinA and encircled all the nuclei, whereas torsinA fully encircled only a subset of nuclei. This pattern of PDI staining was similarly observed with another ER marker (ER-Tracker Green) in our previous study of cultured glial cells (Iwabuchi et al., 2014b).

### 3.3. Endogenous WT-torsinA is present mainly near the Golgi apparatus but not in the ER

We further analyzed the subcellular localization of endogenous torsinA in the rat glial cells. The main, paranuclear immunoreactivity of torsinA was distinct from the distribution of the ER (Fig. 2), suggesting that it was localized in a non-ER organelle such as the Golgi apparatus. In order to test this hypothesis, the cultured rat hippocampal glial cells were stained by double-immunocytochemistry for torsinA and the *cis*-Golgi marker GM130 (Horton et al., 2005) (Fig. 3). They were imaged by confocal microscopy to provide high spatial resolution. In two representative fields (upper and lower rows of (Fig. 3A), the paranuclear torsinA signals were co-localized extensively with GM130 signals. They were present near all nuclei, whose positions were indicated by non-specific nucleoplasmic staining for torsinA. A very weak, diffuse torsinA signal was observed in the cytoplasm. The field in the top-right panel is shown at different heights in a z-stack (Fig. 4). The co-localization of torsinA and GM130 signals was maintained in all images.

**Fig. 3.**
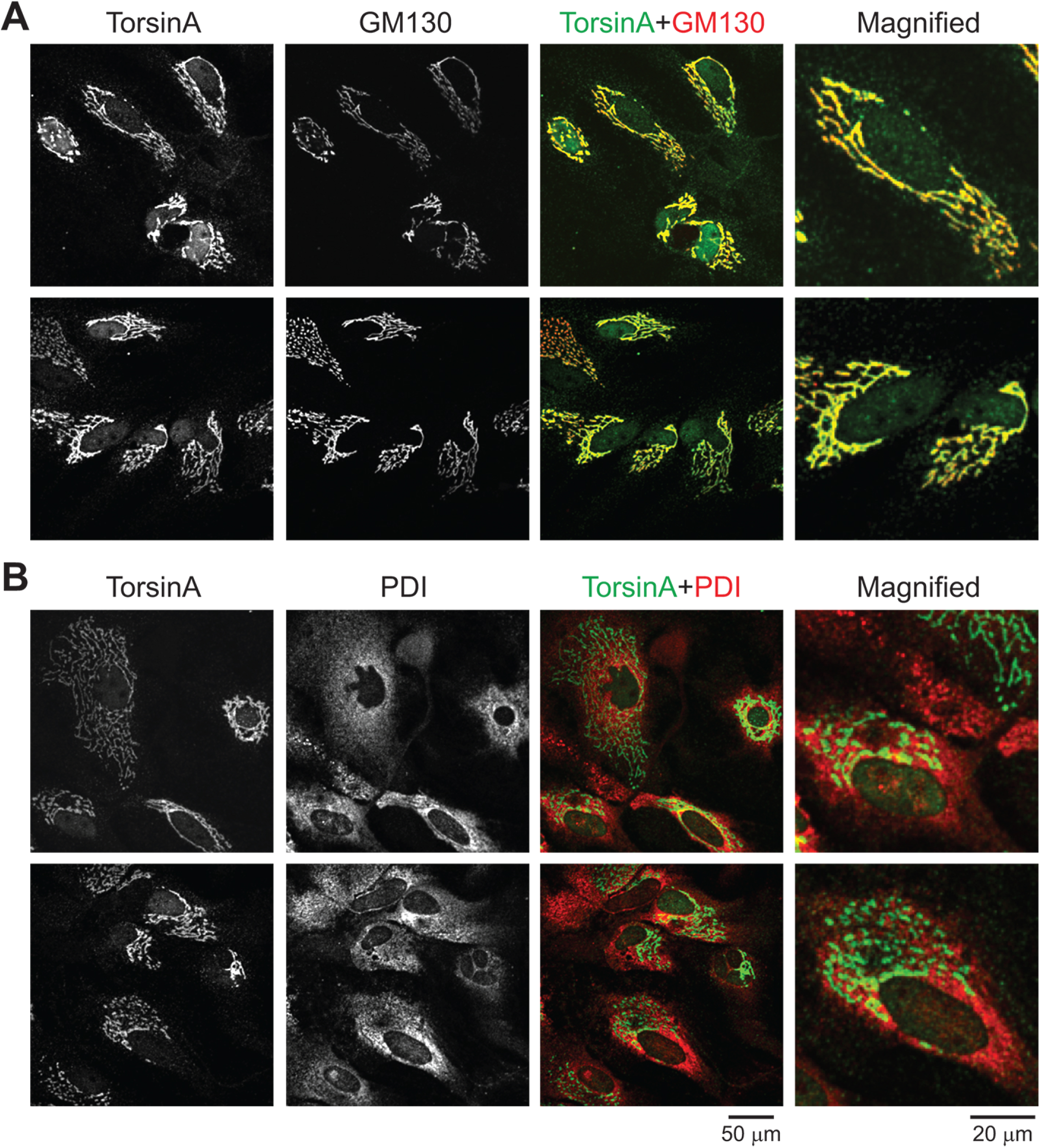
Localization of endogenous WT-torsinA relative to the Golgi and ER in cultured glial cells from the rat hippocampus. **A**) Cells double-stained for torsinA (green channel) and the Golgi marker GM130 (red channel). The two signals are extensively co-localized. TorsinA channel has additional, weak nucleoplasmic staining as well as weaker cytoplasmic staining. **B**) Cells double-stained for torsinA and PDI. The two distributions are similar but not identical. Two representative fields are shown in both **A** and **B**. TorsinA was visualized using the Alexa Fluor 488 dye, and GM130 or PDI using the Alexa Fluor 568 dye, by confocal microscopy.

**Fig. 4.**
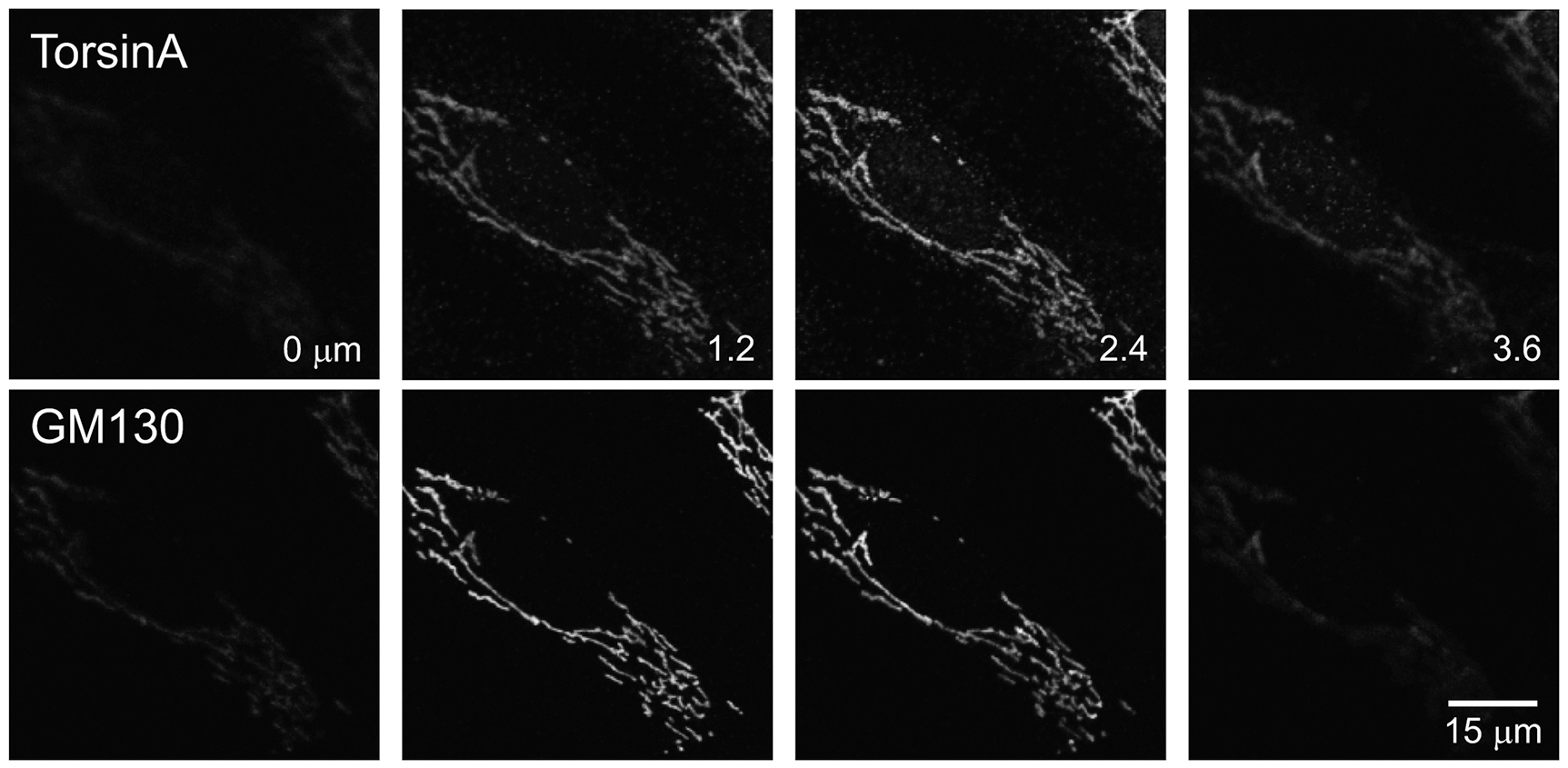
Co-localization of endogenous WT-torsinA and Golgi signals in cultured glial cells from the rat hippocampus. Confocal images of the same field as in Fig. 3A. Images are shown at different heights in a z-stack, after 3 consecutive images were averaged and filtered with a Gaussian function. The heights in μm represent the vertical distances between the middle of 3 images.

In sharp contrast, the torsinA and PDI signals were not co-localized (Fig. 3B). The main torsinA signal was confined to narrower spaces than that for PDI, consistent with the results obtained using widefield optics (Fig. 2B). A very weak and diffuse torsinA signal was present in the cytoplasm, and seemed to be co-localized with PDI.

### 3.4. Endogenous WT-torsinA is not detectable in the nuclear envelope

Since overexpressed WT-torsinA is localized strongly in the ER but also weakly in the nuclear envelope (Goodchild and Dauer, 2004; Giles et al., 2008), we stained the cells for torsinA and a marker of the nuclear envelope, NPC protein (Fig. 5A). Confocal imaging showed that these proteins were not co-localized (Fig. 5A). NPC staining was observed at the periphery of individual nuclei of the glial cells, where the nuclear envelope signal was located. It appeared to be present also in the nucleoplasm, though at a slightly weaker intensity. However, in cultured neurons which have thicker bodies than the glial cells, nucleoplasmic staining by NPC was absent (i.e. staining was limited to ring-like structures; data not shown). These data suggest that the seemingly nucleoplasmic NPC signal in glial cells stems from nuclear envelope that lies horizontally below or above the nucleoplasmic section, due to poor optical resolution in the z direction in comparison to x-y direction (Egner and Hell, 2006). The fact that torsinA was not co-localized with NPC at the nuclear periphery indicates that torsinA signal was not present in the nuclear envelope at a detectable level.

**Fig. 5.**
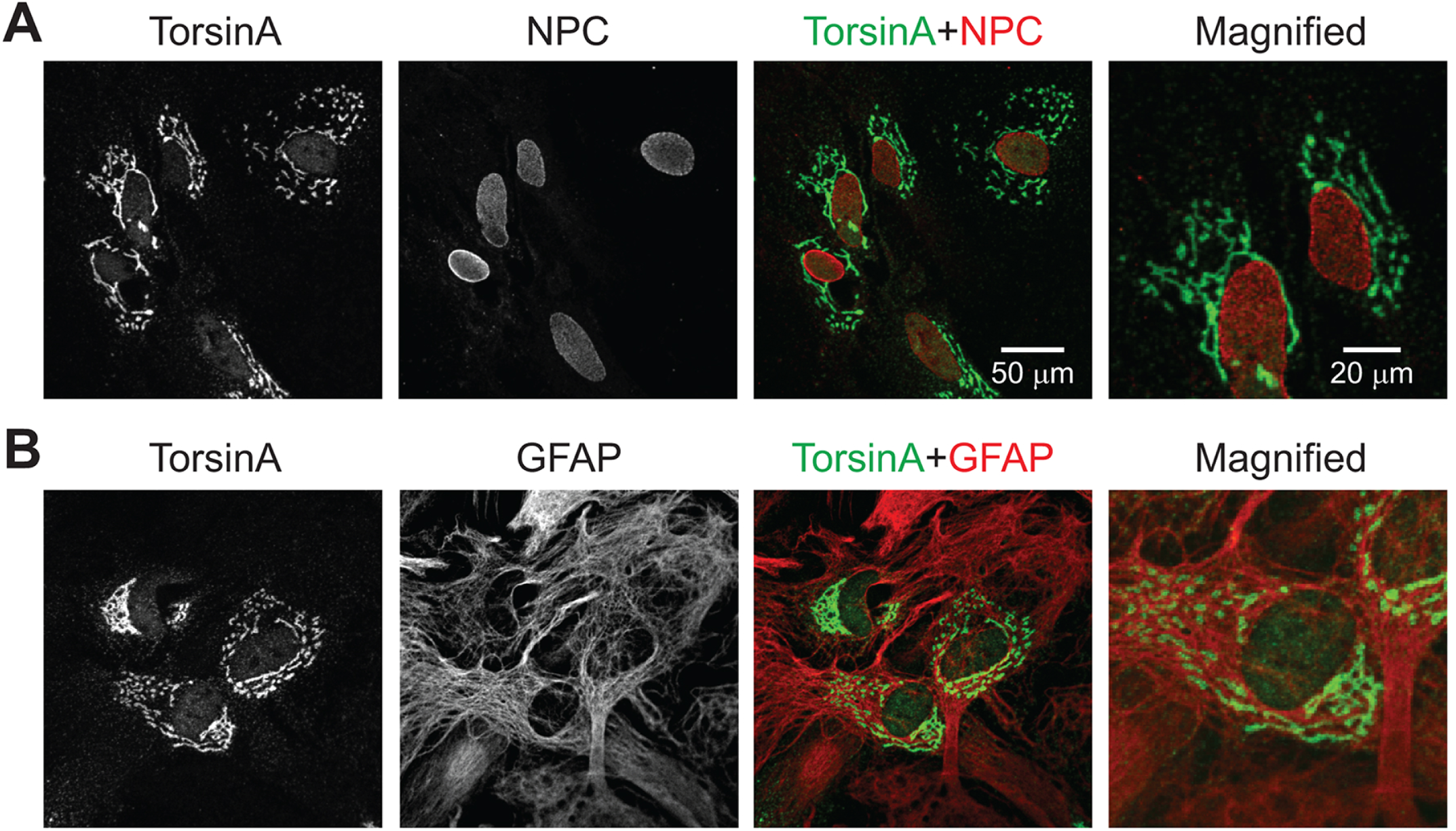
Localization of endogenous WT-torsinA relative to that of nuclear envelope and cytoskeleton in cultured glial cells from the rat hippocampus. **A**) Cells double-stained for endogenous torsinA and the nuclear envelope marker, nuclear pore complex (NPC) protein. **B**) Cells double-stained for endogenous torsinA and GFAP. TorsinA was visualized using Alexa Fluor 488 dye (green channel), and NPC or GFAP using Alexa Fluor 568 dye (red channel), by confocal microscopy. In both **A** and **B**, the distribution of torsinA was different from that of the non-Golgi markers.

Since overexpressed WT-torsinA is co-localized with GFAP in human astrocytes (Armata et al., 2008), we also double-stained the cells for endogenous torsinA and GFAP (Fig. 5B). Confocal microscopy showed that the distributions of these proteins were also distinct. The GFAP staining occupied almost the whole image field, whereas the main torsinA staining was paranuclear and very limited in spatial extent in comparison to that of GFAP.

These results demonstrate that the signals of the endogenous torsinA in WT rat glial cells (Figs. 2-5) were distributed differently from those of the overexpressed WT-torsinA (Fig. 1).

### 3.5. Distributions of markers of Golgi, ER and cytoskeleton in glial cells

Our experiments revealed considerable differences in the distributions of the Golgi apparatus (GM130; Fig. 3A), ER (PDI; Figs. 2B, 3B) and cytoskeleton (GFAP; Figs. 1A, 5B). In terms of stained area, they were in the order of Golgi < ER << cytoskeleton. This result was against a simple assumption that ER and cytoskeleton are distributed similarly within cells. In the next three figures (Figs. 6-8), we further analyzed the basic morphological features of cultured rat glial cells, in order to test whether these differences were due to cell-to-cell variations or other factors which can affect the interpretation of torsinA localization.

**Fig. 6.**
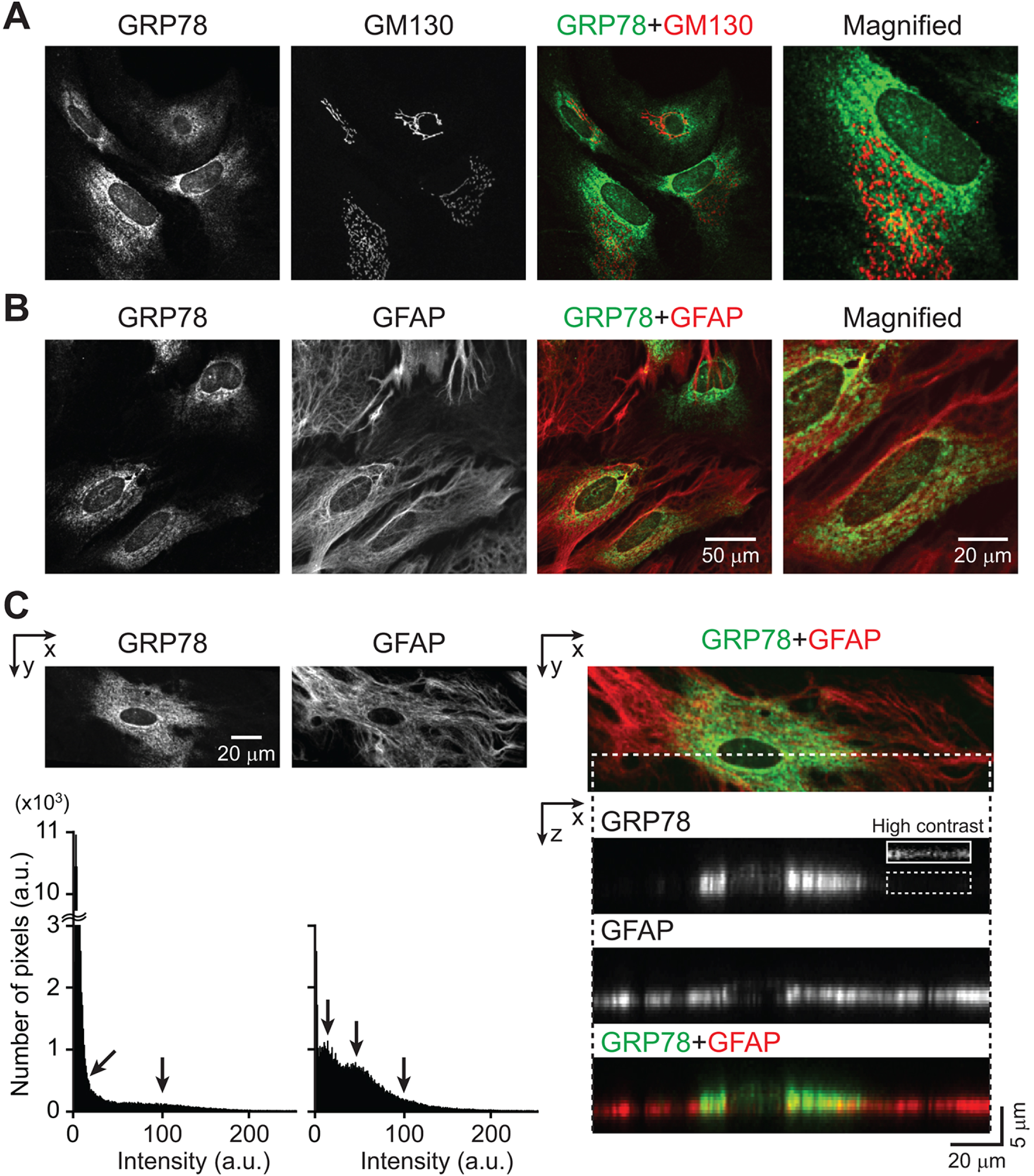
Relationship between Golgi, ER and GFAP in cultured glial cells from the rat hippocampus. **A**) Cells double-stained for the ER marker GRP78 and the Golgi marker GM130. **B**) Cells double-stained for GRP78 and the astrocyte cytoskeleton GFAP. **C, left**) GRP78 and GFAP staining (x-y images) and the corresponding intensity histograms. Image contrast was adjusted such that the strongest and weakest signals in the same image were assigned 255 and 0 values on an 8-bit scale. Arrows in the histograms represent positive signals. Tallest peak near the 0-intensity level in each histogram represents background noise (not labeled). **C, right**) GRP78 and GFAP images are overlaid (x-y image, top). The x-z images (3 panels) were obtained through the dotted line in the x-y image. Coverslip lies at the bottom of the x-z images where no signals are found. Image contrast was re-adjusted such that the strongest and weakest signals in the same x-z image were assigned 255 and 0 values on an 8-bit scale. Dotted rectangle in the GRP78 image shows a region distal to the nucleus. This region is shown with an enhanced image contrast in the continuous rectangle, demonstrating the presence of weak GRP78 signal. GRP78 was visualized using Alexa Fluor 488 dye (green channel), and GM130 and GFAP were visualized using Alexa Fluor 568 dye (red channel), by confocal microscopy. These results suggest that GRP78 signal in the x-y plane reflects the variable thickness of the cell. ER was present where GFAP was localized, but was not easy to detect away from nuclei because the image contrast was adjusted to detect the brightest signal (near the nuclei). In the x-y plane, GFAP signal appeared to be universally present and was of uniform intensity.

#### Golgi-ER relationship

In double-staining experiments, the Golgi was labeled by GM130 and the ER was labeled by the marker GRP78/BiP (Fig. 6A). The GM130 signal was discrete. In contrast, GRP78 signal was diffuse, similar to the PDI signal (Figs. 2B, 3B). Overlaid images show that GRP78 was distributed more widely than GM130 in all cells (Fig. 6A). GRP78 signal was not detectable throughout the image field, again similarly as PDI signal (Figs. 2B, 3B). This finding raised the possibility that the glial culture in the imaged field was not confluent. However, analysis of other coverslips by DIC and fluorescence imaging showed that GRP78 signal was confined to regions near the nuclei even where glial cells covered all parts of the field (Fig. 7A). We also confirmed that the GM130 signal was paranuclear, fully encircling some nuclei but not in others (Fig. 7A).

**Fig. 7.**
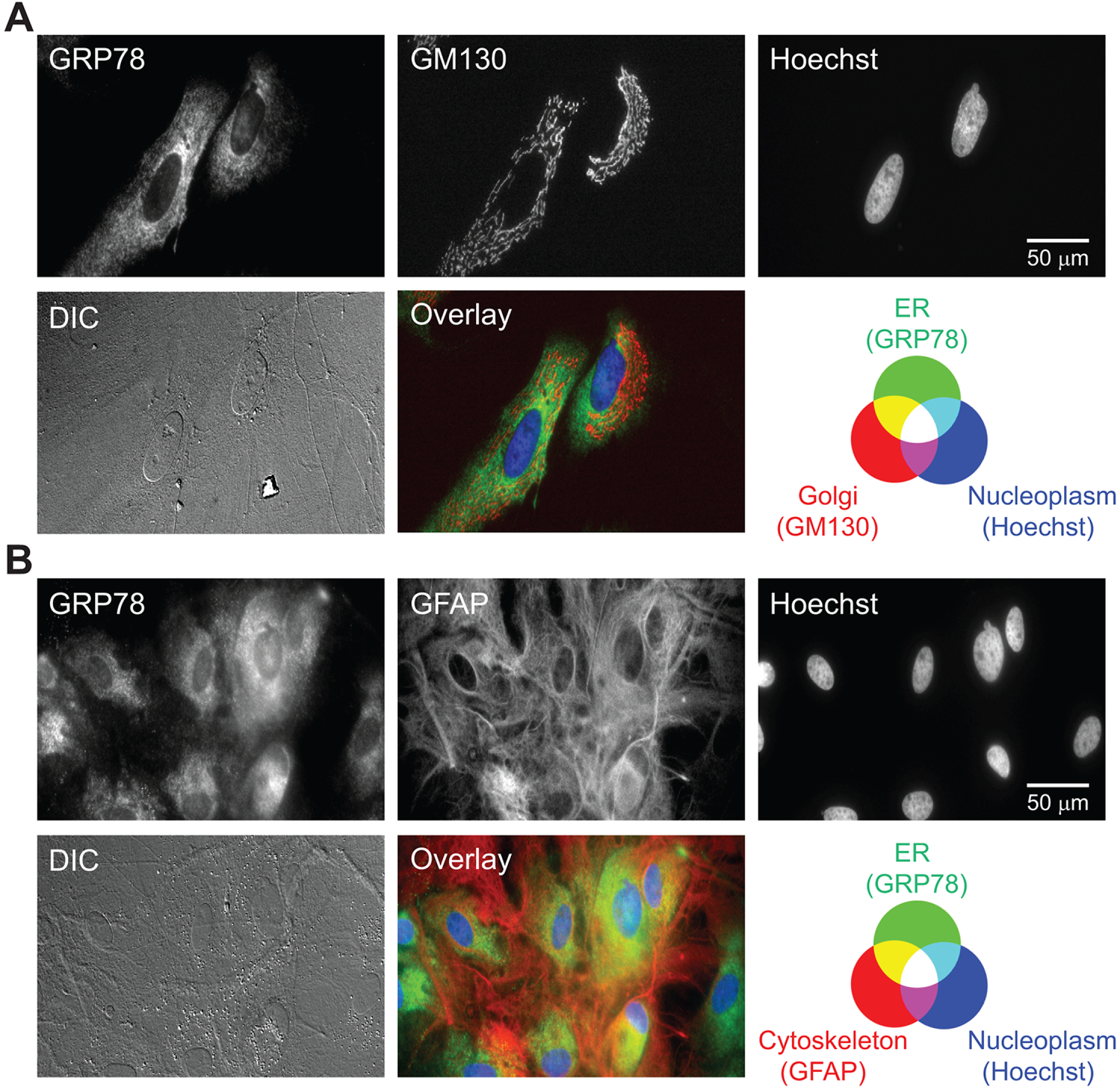
Relationship between Golgi, ER and GFAP in cultured glial cells from the rat hippocampus. **A**) Cells double-stained for the ER marker GRP78 and the Golgi marker GM130, and counterstained for nuclei (Hoechst dye). **B**) Cells double-stained for GRP78 and the astrocyte cytoskeleton marker GFAP, and counterstained for nuclei. In this figure, GRP78 was visualized using Alexa Fluor 488 dye (green channel), and GM130 and GFAP were visualized using Alexa Fluor 568 dye (red channel), by widefield microscopy.

#### ER-cytoskeleton relationship

Double-staining experiments with anti-GRP78 and GFAP antibodies were carried out. Confocal overlays show that GFAP was distributed more widely than GRP78 (Fig. 6B). Another experiment confirmed the differences in the distributions of GRP78 and GFAP (Fig. 7B). The cells formed a confluent layer as judged by DIC image, GFAP staining was nearly uniform, but GRP78 staining was confined to regions closer to the nuclei. Neither GRP78 nor GFAP was present in the nucleoplasm (Fig. 7B). Thus, ER << cytoskeleton in the immunocytochemical images.

The apparent difference (ER << cytoskeleton) in the distributions of ER proteins (PDI and GRP78) and cytoskeletal GFAP in the horizontal (x-y) plane can be explained by the combined effects of four factors that involve distribution in the vertical (x-z) plane.

First, the GRP78 and GFAP signals were present at different focal levels. Confocal images in a z-stack show that the visible GRP78 signal was distributed in a broader thickness level (~0.8 to ~4.4 µm in height) than GFAP (~0.8 to ~2.8 µm in height) (Fig. 8). In addition, the strongest signal was obtained at higher height for GRP78 than GFAP (arrows in Fig. 8). These points were confirmed in a second analysis (Fig. 6C). Specifically, images of another cell show that GFAP was more widely distributed than GRP78 in the x-y plane (top panels of Fig. 6C). The images in the x-z plane show that the GRP78 signal was variable in thickness, with the thickest part immediately surrounding the nucleus. In contrast, GFAP signal was clustered in a thinner layer and located near the bottom of the cell (immediately above the coverslip).

**Fig. 8.**
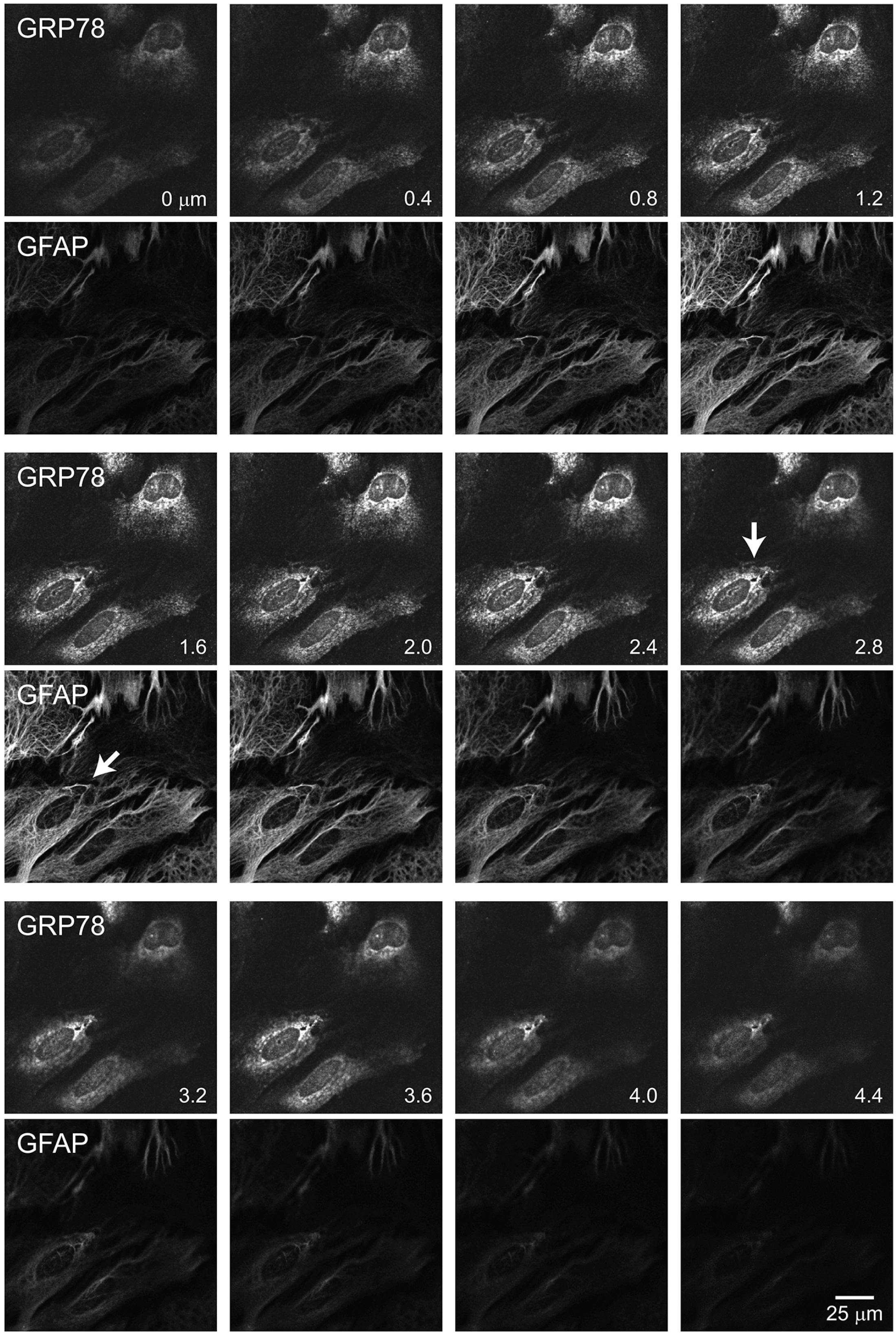
Relationship between the ER and GFAP in cultured glial cells from the rat hippocampus. The same field as shown in Fig. 6B, with optical sections taken starting near the coverslip and moving up along the z axis at 0.4-µm intervals. Arrows point to where in the z stack the GRP78 and GFAP signals are strongest.

Second, GRP78 and GFAP signals showed very different variations in intensity. This point is illustrated by the intensity histograms obtained from the x-y images (bottom-left panels of Fig. 6C). The intensity of the GRP78 signal varied broadly (arrows), especially covering the weak intensity levels (tilted arrow), whereas that of the GFAP signal was relatively uniform and covered intermediate intensity levels (arrows). The x-z images show that the brightest GRP78 signal came from regions surrounding nucleus. Importantly, they also show that GRP78 signal was present far from nucleus, although the intensity at this location was weak (boxed region). In contrast, GFAP signal was of relatively uniform intensity, due partly to its uniform thickness.

Third, intensity of the GRP78 signal near the nucleus is amplified by its thickness. This is due to poor optical resolution in the z-direction (Egner and Hell, 2006). The vertical signal spreads from a point source (point spread function) and can reach >1 µm when viewed by confocal microscopy. Thus the signals from neighboring images can accumulate in the vertical direction, because the interval of image acquisition (voxel height) was 0.40 µm in this series. The additive effect is less for signals that have thinner structures in the x-z plane, e.g. GRP78 signal far from nucleus and GFAP signal.

Lastly, a large intensity variation of the GRP78 signal forced the weaker signals to appear dimmer. This is because the maximum and minimum intensity in each channel were assigned the values of 255 and 0 on an 8-bit scale, and the brighter signal near nuclei capped the intensity distribution. GFAP signals were more uniform in intensity and thus suffered less from the dimming effect of intensity variations.

In summary, ER (GRP78) and cytoskeletal (GFAP) signals were both ubiquitously present. However, the signals appeared as ER << cytoskeleton in terms of area, because the ER signal in the x-y plane reflected the variable thickness of the cell. This analysis further supports the notion that the main signal of endogenous torsinA is paranuclear and in the Golgi apparatus, because its distribution is markedly different from that GFAP, and rather similar to that of the ER, which is prominent near the nucleus where the Golgi is located.

### 3.6. Distribution of endogenous torsinA proteins in mouse glial cells

Thus far, we have analyzed rat glial cells. We next examined the subcellular localization of WT- and ΔE-torsinA proteins in cultured mouse glial cells. These cells were obtained from the hippocampus and striatum of the ΔE-torsinA knock-in mouse model of DYT1 dystonia (Goodchild et al., 2005). GFAP staining revealed that most were astrocytes (Fig. 9).

**Fig. 9.**
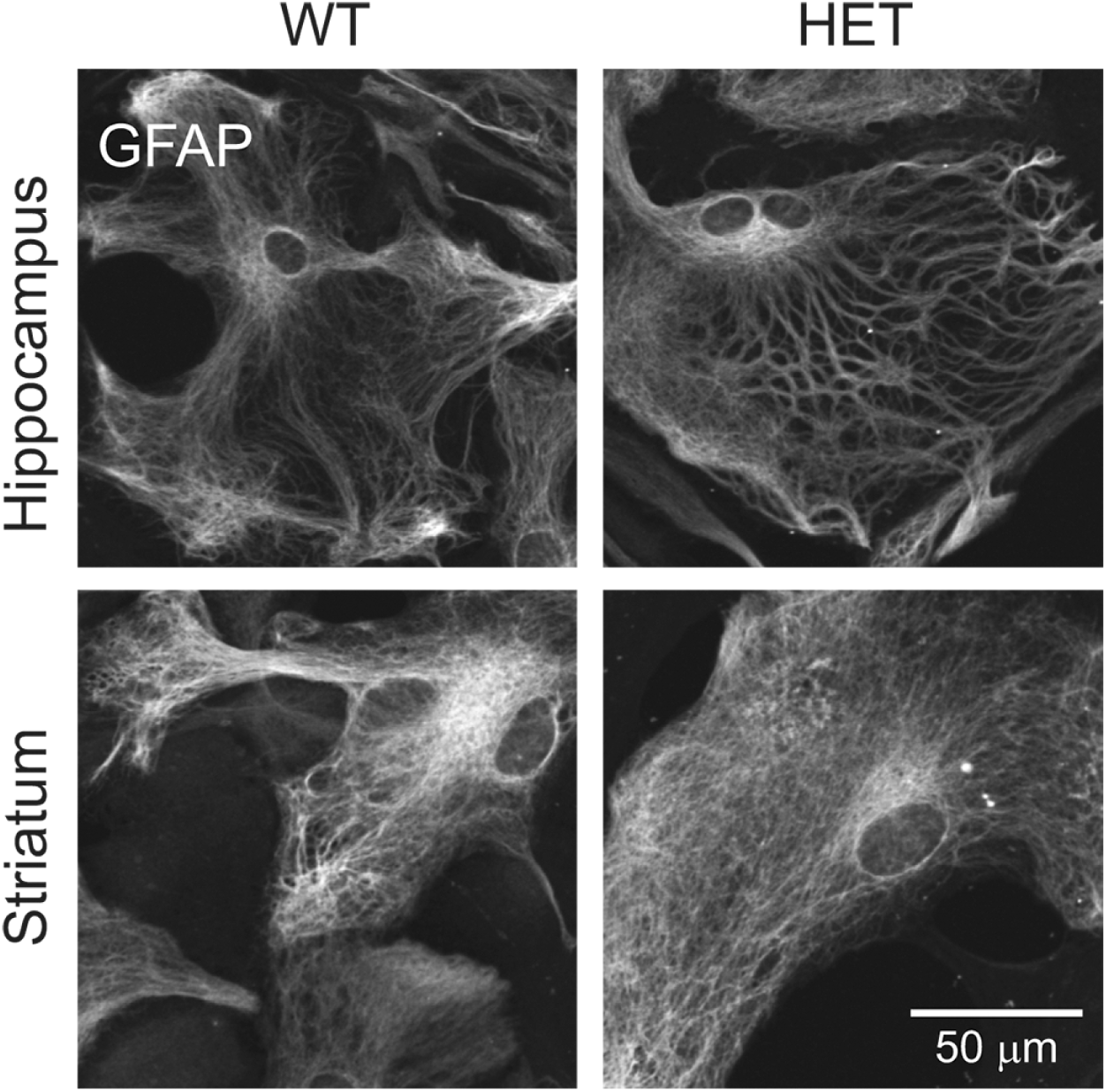
GFAP staining of cultured glial cells from the mouse hippocampus and striatum. Cells from the two different brain regions of WT and heterozygous (HET) ΔE-torsinA knock-in mice. GFAP was visualized using the Alexa Fluor 488 dye (green channel), by confocal microscopy.

The striatum is the brain region that has been implicated in the pathophysiology of dystonia. In striatal cells from the WT mouse, distribution of the immunoreactive torsinA (top panels of Fig. 10A,B) was similar to that in WT rat hippocampal cells (Fig. 3). In striatal cells from the HET mice, its distribution was also the same (bottom panels, Fig. 10A,B). The main torsinA signal was co-localized with GM130 signal in the striatal glial cultures obtained from both WT and HET mice (Fig. 10A). A minor component of the signal was present in the nucleoplasm but not in the putative nucleoli. A weaker, diffuse signal was present in the cytoplasm. The main torsinA signal was not co-localized with PDI (Fig. 10B). In the torsinA channels, we observed neither cytoplasmic inclusion bodies nor perinuclear staining suggestive of nuclear envelope staining, as opposed to when ΔE-torsinA was overexpressed. Findings were the same for the mouse hippocampal glial cultures (data not shown). Thus the distribution patterns of endogenous torsinA were identical in the glial cells of WT and HET mice, and were the same as in WT rat hippocampus (Fig. 3A).

**Fig. 10.**
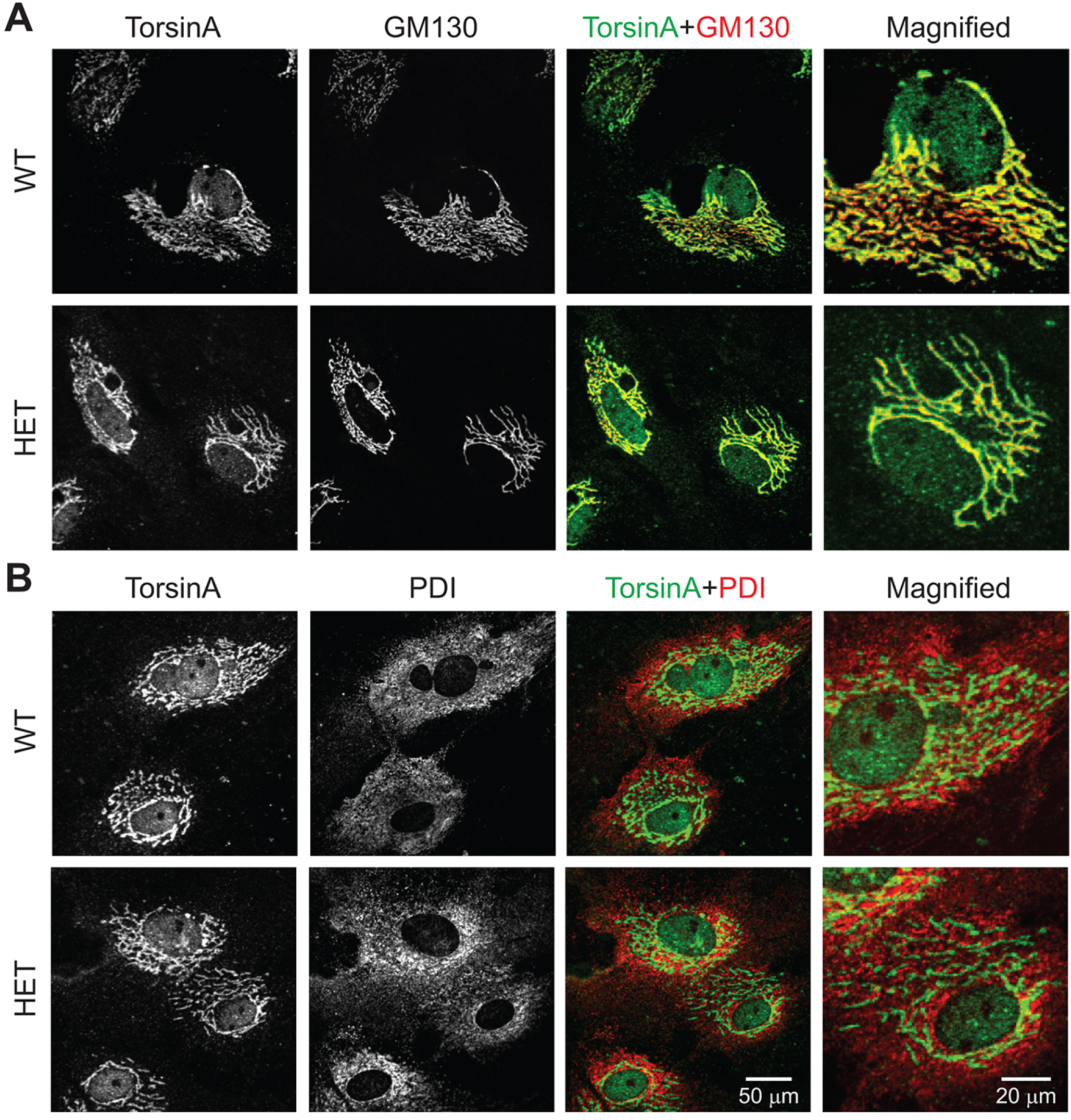
Localization of endogenous WT-torsinA in cultured glial cells from the mouse striatum. **A**) WT and HET glial cells double-stained for endogenous torsinA and GM130. The main torsinA and GM130 signals are extensively co-localized, although torsinA is additionally present at lower levels in the nucleoplasm and cytoplasm. **B**) WT and HET cells double-stained for endogenous torsinA and the ER marker PDI. The two distributions are similar but not identical. TorsinA was visualized using the Alexa Fluor 488 dye (green channel), and GM130 and PDI were visualized using the Alexa Fluor 568 dye (red channel), by confocal microscopy.

### 3.7. Quantitative analysis of torsinA distribution at endogenous expression level

In order to quantify the data obtained so far, we measured the intensity of torsinA staining in various subcellular compartments. The torsinA intensity in the Golgi apparatus was measured by assigning ROI’s on the GM130 image, and transferring these ROI’s to the torsinA image of the co-stained specimen (Fig. 11A). The intensities of torsinA in the nucleoplasm (excluding nucleoli) and the cytoplasm (ER excluding Golgi) were measured by assigning ROI’s on the PDI image, and then transferring them to the torsinA image of the co-stained specimen (Fig. 11A).

**Fig. 11.**
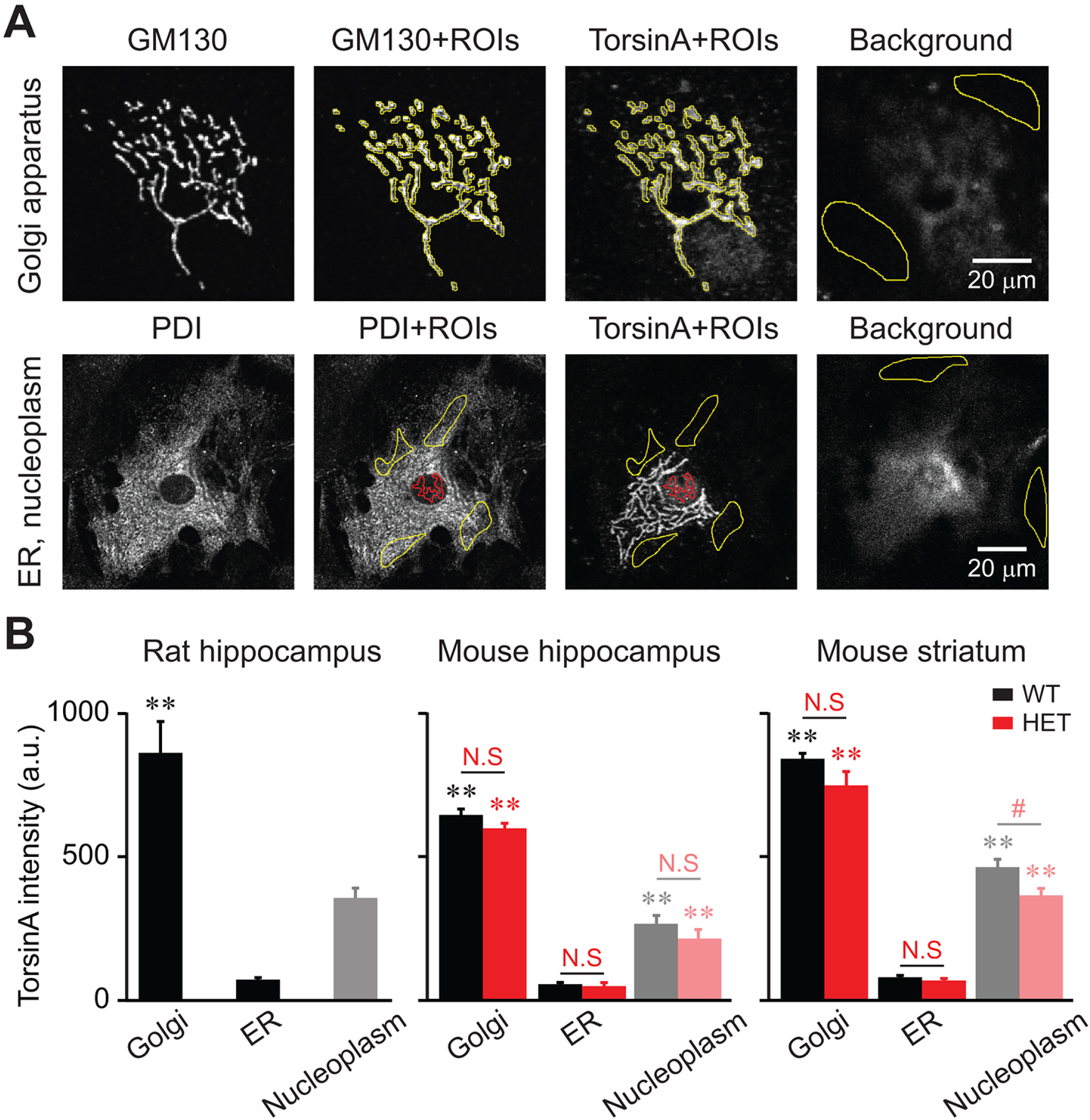
Levels of immunoreactive endogenous torsinA in various subcellular compartments. **A**) Method for measuring torsinA intensity in the Golgi, ER and nucleoplasm. Cells were co-stained for torsinA and GM130 (top), or for torsinA and PDI (bottom). The Golgi apparatus was measured by assigning regions of interest (ROI’s) in the GM130 channel, and transferring them to the torsinA channel. The pixel intensities in the ROI’s were averaged to give a single value for one cell. The signal in the ER and nucleoplasm was measured by assigning the ROI’s in the PDI channel, and transferring them to the torsinA channel. The pixel intensities in each compartment were then averaged, providing a single value each for one cell. These values were used for numerical analysis, after subtracting the averaged background intensity. The background regions were assigned as the dimmest regions for the highest image in a z-stack (i.e. non-cellular region in an image furthest from coverslip level). The nucleoplasmic measurement excluded nucleoli. The cytoplasmic measurement excluded Golgi. TorsinA was visualized using the Alexa Fluor 488 dye (green channel), and GM130 and PDI were visualized using the Alexa Fluor 568 dye (red channel), by confocal microscopy. **B**) Quantitation of endogenous torsinA in glial cells, cultured from the WT rat hippocampus (left panel), mouse hippocampus (middle panel), and mouse striatum (right panel). In each panel, the y axis represents absolute pixel intensity after the background intensity was subtracted. Imaging conditions were the same for all samples within each panel, allowing statistical comparisons of staining intensity. The conditions differed for each panel. In all 5 cases examined, the intensity of Golgi staining was significantly higher than that for the ER (asterisks indicate significance): p<10^−4^, n = 25, 29 cells in WT rat hippocampus (left); p<10^−19^, n = 13, 14 cells in WT mouse hippocampus (middle); p<10^−14^, n = 9 cells in HET mouse hippocampus (middle); p<10^−7^, n = 11, 12 cells in WT mouse striatum (right); p<10^−14^, n = 16, 17 cells in HET mouse striatum (right). There were no genotypic differences in torsinA intensity (NS, heterozygous vs. WT): p>0.05 in Golgi of mouse hippocampus (middle); p>0.1 in ER of mouse hippocampus (middle); p>0.3 in Golgi of mouse striatum (right); p>0.1 in ER of mouse striatum (right). Nucleoplasmic staining was not analyzed in detail (gray), because it was considered non-specific.

TorsinA intensity was measured in the glial cells of the hippocampus of WT rats, the hippocampus of WT and HET ΔE-torsinA knock-in mice, and the striatum of WT and HET ΔE-torsinA knock-in mice (Fig. 11B). In all five cases, torsinA staining in the Golgi apparatus was significantly more intense than that in the ER (p<10^−4^; n = 9-29 cells per measurement). When normalized by the averaged intensity in the ER, the Golgi intensity (Golgi/ER) in the five cases ranged from ~11.3 (WT mouse hippocampus) to ~13.8 (HET mouse hippocampus).

In addition, there was no significant genotypic difference in the Golgi or ER signals in either the hippocampal (HET vs. WT, p>0.05, n = 9 cells) or striatal (p>0.1, n = 17 cells) mouse glial cells.

Thus, the ER was not the main site of localization of endogenous WT-torsinA. Furthermore the distribution of ΔE-torsinA did not differ from that of the WT protein.

### 3.8. Relative expression levels of endogenous torsinA in rat glial cells and neurons

Although glial cells have been shown to express torsinA (Kim et al., 2010; Puglisi et al., 2013), glial expression was not detected in some studies (Shashidharan et al., 2000; Rostasy et al., 2003; Martella et al., 2009). In order to find the reason for the discrepancy, we evaluated the glial expression level relative to the neuronal expression. Neuron-glial co-cultures were established from the rat hippocampus and were co-stained for torsinA and GFAP (Fig. 12A). Glial torsinA was defined as the signal nearest the glial nuclei, that were negative for GFAP staining. The remaining torsinA staining was found to be neuronal in origin (manuscript in preparation). It was possible to distinguish between glial and neuronal torsinA signals also because the glial cells were flatter (a few µm thick, Fig. 6C) than the neurons (>10 µm thick; data not shown). This leads to glial torsinA signals being located closer to the coverslip than neuronal torsinA (Fig. 12A).

**Fig. 12.**
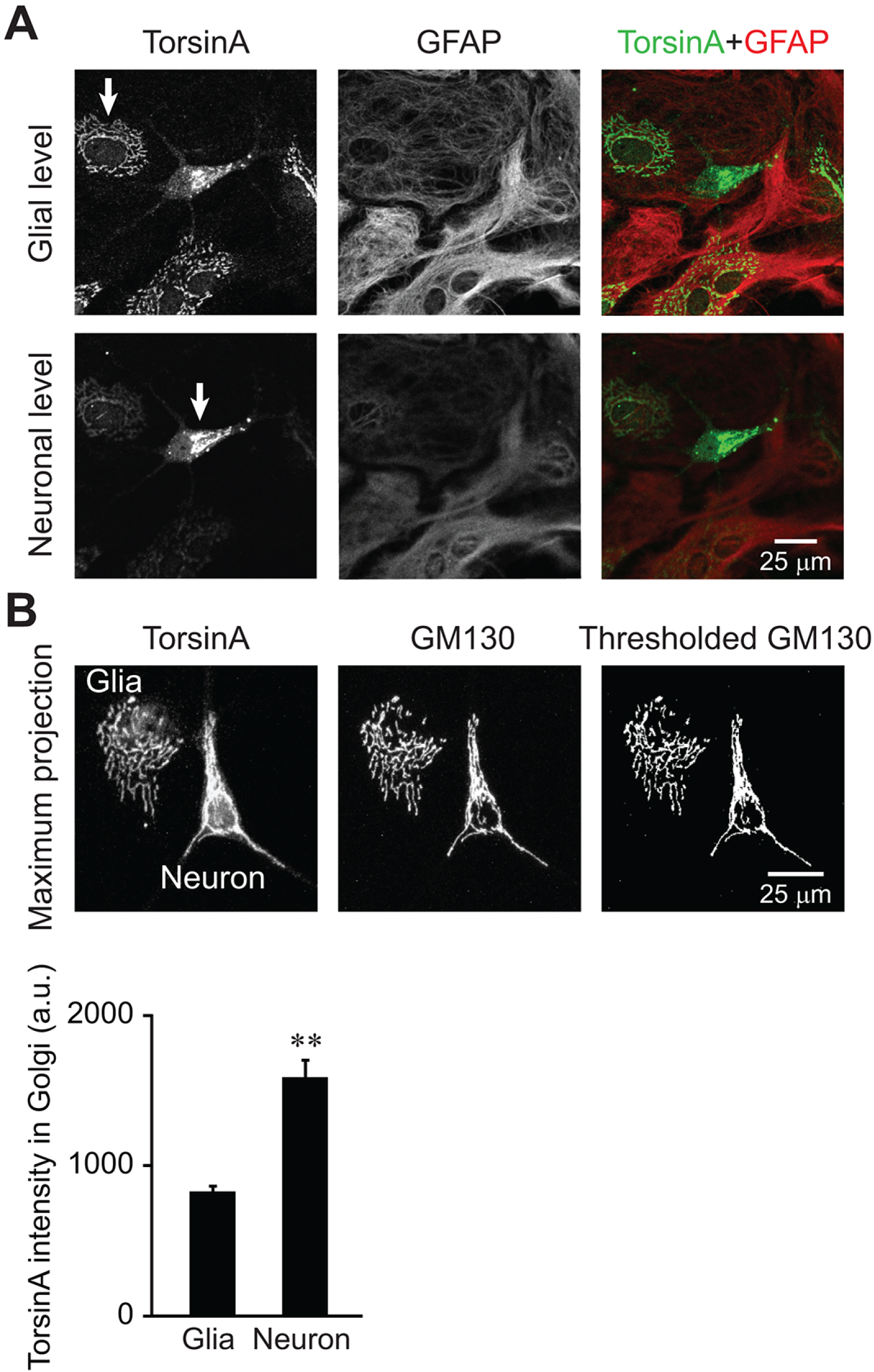
Levels of torsinA in glial cells and neurons. Neuron-glial cultures were prepared from the hippocampus of WT rats. **A**) Co-staining for torsinA and GFAP. The glial torsinA signal (arrow, glial level) was found in the same plane as GFAP, whereas the neuronal torsinA signal (arrow, neuronal level) was further from the coverslip. **B**) Method for measuring torsinA intensity in the glial cells and neurons. It was the same as in Fig. 8A. The graph shows the torsinA intensity, after background intensity in the non-cellular region was subtracted (as in Fig. 8A). The intensity of Golgi staining was significantly higher in neurons than in glial cells, with the average intensities of 1564 and 855 arbitrary units, respectively (p<10^−4^, n = 7 neuron-glia pairs).

The images in a confocal z-series were compressed using a maximum intensity projection (Fig. 12B). This algorithm identifies the maximal intensity at all focal levels at a single pixel (i.e. maximal intensity at different z coordinates for a single x-y coordinate pair). The GM130 staining marked the Golgi, and ROI’s based on this staining were transferred to the torsinA images (Fig. 12B). The averaged pixel intensity in the Golgi apparatus of each cell was significantly higher in neurons than in glial cells (p<10^−4^; n = 7 glial cells and neurons) (Fig. 12C). Its neuron/glia ratio was ~1.8. The low level of glial signal could have led to the omission of glial expression in previous studies.

## 4. DISCUSSION

Using rodent glial cells in primary culture, we have demonstrated that: 1) overexpressed torsinA proteins were distributed as predicted by the mis-localization model, 2) the signals of endogenous torsinA were detectable near the Golgi apparatus, 3) localization of endogenous torsinA in the *Tor1a*^+/ΔE^ mice (which express one WT and one ΔE allele) did not differ from that of the WT-torsinA, and 4) the endogenous distribution was the same regardless of brain region and rodent species examined. This study demonstrates a discrepancy between the localization of overexpressed and endogenous torsinA proteins in a single series of experiments.

### 4.1. Exogenous expression of torsinA in glial cells

TorsinA overexpression has been reported in the human glia-derived tumor (glioma) line Gli36 cells (Bragg et al., 2004a; Bragg et al., 2004b). The overexpressed WT-torsinA was diffuse throughout the cytoplasm, co-localized with PDI and was found around the nucleus. In contrast, the overexpressed ΔE-torsinA led to the formation of cytoplasmic inclusions and perinuclear staining that was co-localized with nuclear envelope markers laminB and nucleoporin (Bragg et al., 2004a; Bragg et al., 2004b). With respect to astrocytes, the outcome of torsinA overexpression had been reported in only one study. Human WT-torsinA had been overexpressed in cultured human astrocytes of a non-specified brain region (Armata et al., 2008). The signal was diffuse throughout the cytoplasm and co-localized with GFAP. The overexpression of ΔE-torsinA has not been reported in any type of astrocytes. We found that overexpressed WT-torsinA in rat astrocytes results in diffuse cytoplasmic staining and that overexpressed ΔE-torsinA results in staining of inclusion bodies (Fig. 1). Our observation is consistent with the findings from glioma cells and human astrocytes. This was also consistent with the overexpression results from neurons (Nery et al., 2011), neural tumor cells (Hewett et al., 2000; Gonzalez-Alegre and Paulson, 2004; Misbahuddin et al., 2005; Gordon and Gonzalez-Alegre, 2008), and non-neuronal, non-glial cells (Kustedjo et al., 2000; Goodchild and Dauer, 2004; Vander Heyden et al., 2009). Thus our data support the mis-localization model of ΔE-torsinA when the protein is overexpressed (Harata, 2014).

### 4.2. Endogenous expression of torsinA in glial cells

Astrocytes and glia-related cells (such as glioma cells) have been reported to express endogenous, wild-type torsinA. Western blot analysis revealed endogenous torsinA protein in the mouse glioma line GL261 cells (Hewett et al., 2000), as well as in cultured cerebral cortical astrocytes isolated from WT mice (Goodchild et al., 2005; Kim et al., 2010). In addition, immunohistochemistry has revealed the expression of endogenous torsinA in astrocytes in the rat hippocampus *in vivo* (Zhao et al., 2008), in the Bergmann glial cells and bushy astrocytes of the mouse cerebellar cortex *in vivo* (Puglisi et al., 2013), and in glial cells of neuron-glia cultures (Koh et al., 2013). In contrast, immunohistochemistry studies did not detect endogenous protein in glial cells from the brains of human control subjects (Shashidharan et al., 2000; Konakova et al., 2001; Rostasy et al., 2003; Siegert et al., 2005), of DYT1 dystonia patients (Rostasy et al., 2003; McNaught et al., 2004), or of WT mice (Martella et al., 2009). Similarly, *in situ* hybridization did not detect the torsinA mRNA in glial cells of the human brain, although it was present in neurons (Augood et al., 1999). Notably, the lack of detectable mRNA signal did not necessarily reflect a lack of this specific mRNA, because the detection sensitivity in these experiments may have been low (Augood et al., 1999).

We propose that the above discrepancy is due to differences in the expression levels of torsinA in astrocytes and neurons. The neuronal expression is ~1.8-fold higher than astrocytes in a projected plane of each cell (Fig. 12). Note that this value represents the pixel intensity within a single plane and that the total expression level per neuron will reflect the total signal integrated over the cell volume, and thus is expected to be much higher. In previous studies that detected glial torsinA, the glial cells were evaluated under one of the following sets of conditions that favor feasible detection of glial signals: 1) in a preparation devoid of neurons, i.e. in pure glial cultures (Hewett et al., 2000; Goodchild et al., 2005; Kim et al., 2010); 2) under a cellular stress that upregulated glial torsinA, i.e. in the rat brain after ischemia (Zhao et al., 2008); 3) using sensitive confocal or widefield fluorescence microscopy, in mouse brains (Puglisi et al., 2013) or neuron-glia cultures (Koh et al., 2013). In contrast, in the previous imaging studies where glial torsinA was not detectable, the preparations invariably included neurons: human brains (Augood et al., 1999; Shashidharan et al., 2000; Konakova et al., 2001; Rostasy et al., 2003; McNaught et al., 2004; Siegert et al., 2005) and mouse brains (Martella et al., 2009). In those cases, the imaging conditions could have been optimized for visualizing the strong neuronal signals, and the weaker glial signals could have been masked and mistakenly interpreted as negative staining.

Another factor that may have contributed to this discrepancy about whether glial signal is detectable is the narrow, spatial distribution of endogenous torsinA in glial cells. It occupied a small fraction of the glial cells identified by DIC (Fig. 2). Also the glial signal was much less widely distributed than GFAP (Figs. 5B, 12A). Thus, when GFAP is used in search of glial torsinA, it will not be a sufficient guide.

### 4.3. Subcellular distribution of endogenous torsinA in glial cells

Even in studies that have detected endogenous torsinA in the glial cells of WT animal brains (Zhao et al., 2008; Puglisi et al., 2013), the subcellular distribution was not studied in detail. Importantly, however, the spatial domain of torsinA staining appeared to be a small subset of the area delineated by other cytoplasmic markers. For example, in reactive astrocytes of the rat hippocampus *in vivo*, seven days after ischemia, there was upregulation of both torsinA (mRNA and protein levels) and GFAP (protein level). However, torsinA protein was localized to a subcellular region near nucleus, and absent from the extended processes that were positive for GFAP (Zhao et al., 2008). Unfortunately, this study did not assess astrocyte staining prior to ischemia. In another example, in bushy astrocytes of the mouse cerebellar cortex *in vivo*, torsinA signal was strong within the cell bodies and weaker in the GFAP-positive processes (Puglisi et al., 2013). Although these subcellular features were not discussed in the original reports, they are consistent with our findings that torsinA signal was located near nuclei and occupies only a small portion of the GFAP-stained area (Figs. 5B, 12A).

In evaluating the subcellular distribution of torsinA, it is useful to consider morphological relationship of Golgi, ER and cytoskeletal proteins in astrocytes (Figs. 6-8). Partial overlap between torsinA and ER signals is not a guarantee that torsinA is localized to the ER (Figs. 2B, 3B, 10B). This is because Golgi apparatus, which is a distinct structure from ER, can appear colocalized with the ER signal (Figs. 6A, 7A). This situation is especially true in cells where the easily detectable ER signals are confined to the immediate neighborhood of the nucleus. Under such circumstances, the Golgi apparatus becomes confined to a similar subcellular region and can appear to co-localize with the ER signal. Furthermore, a limited distribution of torsinA with respect to GFAP (Figs. 5B, 12A), as with Golgi and ER proteins (Figs. 6B, 6C, 7B, 8), suggests that torsinA localization is not strongly influenced by an interaction with the cytoskeletal protein. Overall, our results indicate that it will be informative to include the Golgi apparatus as a marker in narrowing down the endogenous torsinA localization.

### 4.4. Implications of the current study

We have found markedly different signal distributions of exogenously overexpressed and endogenous torsinA proteins. The reason for this difference is not yet clear. Admittedly, a limitation of the current study is its reliance on cultured cells plated at low density. Although this has significant advantages, including the high spatial resolution and high detection sensitivity, it is possible that the culture conditions influenced the cellular environment and affected the distribution of endogenous torsinA. However, the fact that the overexpression experiments in the current study were also performed on cultured glial cells renders this scenario less likely.

A more plausible explanation for the observed discrepancy is that the exogenous proteins can be subject to general side effects of overexpression. Overexpressed proteins can saturate and overload the native binding or anchoring sites that are used to restrict endogenous proteins spatially. Such overexpressed proteins tend to be distributed to the wrong compartments, for example diffusely in the cytoplasm or aggregated in cytoplasmic inclusions (Gibson et al., 2013). A more specific, unwanted effect has been reported for exogenous torsinA. Overexpression of WT-torsinA in WT neurons induced a nuclear envelope abnormality similar to that seen when WT-torsinA is completely absent in homozygous ΔE-torsinA knock-in mice (*Tor1a*^ΔE/ΔE^) or homozygous *Tor1a* knock-out mice (*Tor1a*^−/−^) (Kim et al., 2010). Moreover, overexpression of WT-torsinA not only failed to rescue the nuclear envelope abnormality in *Tor1a*^−/−^ mice, but aggravated it (Kim et al., 2010). A nuclear envelope abnormality was also observed in central neurons of transgenic mice that overexpress WT-torsinA (Grundmann et al., 2007). These results indicate that excessive manipulation of WT-torsinA, whether up- or down-regulation, is deleterious to cells. Further study will be necessary to delineate the mechanisms whereby endogenous and overexpressed torsinA proteins are distributed, and whereby ΔE-torsinA at an endogenous level impacts cellular functions. Nevertheless, our finding highlights a general concern about relying only on overexpression studies in discussing the pathophysiology of DYT1 dystonia.

## ABBREVIATIONS

DIC: differential interference contrast
ER: endoplasmic reticulum
GFAP: glial fibrillary acidic protein
GFP: green fluorescent protein
GM130: Golgi matrix protein of 130 kDa
GRP78/BiP: glucose-regulated protein of 78 kDa / binding immunoglobulin protein
LED: light-emitting diode
MEM: Minimum Essential Medium
NPC: nuclear pore complex
PBS: phosphate-buffered saline
PDI: protein disulfide isomerase
ROI: region of interest
shRNA: short hairpin RNA
ΔE: deletion of a single GAG codon in the *TOR1A* or *Tor1a* gene

## 5. ACKNOWLEDGMENTS

The authors thank Drs. Phyllis Hanson (Washington University) for WT-torsinA-GFP plasmid construct, Richard Roller (University of Iowa) for the ΔE-torsinA-GFP plasmid construct, Amy Lee, Mark Stamnes, Wayne Johnson (University of Iowa) for discussing the immunocytochemical procedures, Chun-Chi Liang and William Dauer (University of Michigan) for discussion about specificity of the torsinA antibody during the early phase of work, and Jin-Young Koh for technical assistance. This work was supported by grants from the Department of Defense (W81XWH-14-1-0301) and the Dystonia Medical Research Foundation (to N.C.H.).

The authors declare no conflicts of interest.

